# V-ATPase is a universal regulator of LC3 associated phagocytosis and non-canonical autophagy

**DOI:** 10.1101/2021.05.20.444917

**Authors:** Kirsty M. Hooper, Elise Jacquin, Taoyingnan Li, Jonathan M. Goodwin, John H. Brumell, Joanne Durgan, Oliver Florey

## Abstract

Non-canonical autophagy is a key cellular pathway in immunity, cancer and neurodegeneration, characterised by <u>C</u>onjugation of <u>A</u>TG8 to endolysosomal <u>S</u>ingle-<u>M</u>embranes (CASM). CASM is activated by engulfment (endocytosis, phagocytosis), agonists (STING, TRPML1) and infection (influenza), dependent on the ATG16L1 WD40-domain, and specifically K490. However, the factor(s) associated with non-canonical ATG16L1 recruitment, and CASM induction, remain unknown. Here, we investigate a role for V-ATPase during non-canonical autophagy. We report that increased V0-V1 engagement is associated with, and sufficient for, CASM activation. Upon V0-V1 binding, V-ATPase directly recruits ATG16L1, via K490, during LC3-associated phagocytosis (LAP), STING- and drug-induced CASM, indicating a common mechanism. Furthermore, during LAP, key molecular players, including NADPH oxidase/ROS, converge on V-ATPase. Finally, we show that LAP is sensitive to *Salmonella* SopF, which disrupts the V-ATPase-ATG16L1 axis, and provide evidence that CASM contributes to the *Salmonella* host response. Together, these data identify V-ATPase as a universal regulator of CASM, and indicate that SopF evolved in part to evade non-canonical autophagy.

## Introduction

Autophagy is a fundamental degradative process required for nutrient recycling, clearance of damaged organelles and pathogen responses. During autophagy, a collection of ATG proteins induce the formation and maturation of autophagosomes, which enwrap target cargo destined for lysosomal degradation (Choi et al., 2013). A subset of ATG proteins also act in parallel to mediate ‘non-canonical autophagy’, a pathway targeting endolysosomal compartments instead (Florey and Overholtzer, 2012; Heckmann et al., 2017). A diverse set of stimuli and cellular processes induce non-canonical autophagy, including stimulation of the TRPML1 calcium channel (Goodwin et al., 2021), activation of the STING pathway (Fischer et al., 2020), and disruption of endolysosomal ion balance, by pharmacological agents, *Helicobacter pylori* VacA toxin, or infection with influenza A virus (IAV) (Fletcher et al., 2018; Florey et al., 2015; Jacquin et al., 2017). Non-canonical autophagy is also associated with endocytic and engulfment processes, including entosis (Florey et al., 2011), LC3-associated endocytosis (LANDO) (Heckmann et al., 2019) and LC3-associated phagocytosis (LAP) (Martinez et al., 2015; Sanjuan et al., 2007), with essential functions in the killing and clearance of pathogens (Gluschko et al., 2018; Hubber et al., 2017; Martinez et al., 2015), antigen presentation (Fletcher et al., 2018; Ma et al., 2012; Romao et al., 2013), clearance of and cytokine responses to apoptotic cells (Cunha et al., 2018; Florey et al., 2011; Martinez et al., 2011; Martinez et al., 2016; Martinez et al., 2015), clearance of accumulated β-amyloid (Heckmann et al., 2020; Heckmann et al., 2019), vision (Kim et al., 2013), response to influenza infection (Fletcher et al., 2018; Wang et al., 2021) and lysosome biogenesis (Goodwin et al., 2021). While the functional importance of non-canonical autophagy is clear, the mechanisms underlying the regulation and function of this pathway remain incompletely understood.

The canonical and non-canonical autophagy pathways share overlapping machineries, but with important differences. The non-canonical pathway is independent of the upstream autophagy regulators ATG9, mTor and the ULK1 initiation complex (ULK1/2, FIP200, ATG13 and ATG101) (Florey et al., 2011; Martinez et al., 2015). However, both pathways harness the core ATG8 lipidation machinery (ATG3, 4, 5, 7, 10, 12, 16L1). During canonical autophagy, ATG8s are conjugated exclusively to phosphatidylethanolamine (PE), at double membrane autophagosomes (Ichimura et al., 2000). In non-canonical autophagy, ATG8s are conjugated instead to single membrane, endolysosomal compartments, such as phagosomes, endosomes or entotic vacuoles; this process is termed <u>C</u>onjugation of <u>A</u>TG8 to <u>S</u>ingle <u>M</u>embranes (CASM) and involves alternative conjugation to both phosphatidylserine (PS) and PE (Durgan et al., 2021).

ATG16L1 acts as a critical molecular hub, directing canonical and non-canonical autophagy at different sites, via different domains. During canonical autophagy, ATG16L1 is recruited to forming autophagosomes though its coiled coil domain (CCD), which interacts with WIPI2 and FIP200, thereby specifying the site of ATG8 conjugation (Dooley et al., 2014; Gammoh et al., 2013). During non-canonical autophagy, ATG16L1 is recruited instead to preformed endolysosomal membranes to drive CASM. The precise molecular mechanisms underlying this alternative recruitment remain unclear, but the ATG16L1 WD40 C-terminal domain (CTD), and certain ATG16L1 lipid binding motifs, are indispensable (Fletcher et al., 2018; Lystad et al., 2019; Rai et al., 2019). A single point mutation in ATG16L1, K490A, renders cells competent for canonical autophagy, but deficient in non-canonical autophagy, providing a highly specific genetic system to dissect these closely related pathways (Fletcher et al., 2018; Goodwin et al., 2021; Lystad et al., 2019).

The molecular mechanisms of non-canonical autophagy are best studied in the context of LC3-associated phagocytosis (LAP). LAP is dependent on Rubicon/Vps34, which mediates PI3P formation, and NADPH oxidase, which drives the generation of reactive oxygen species (ROS) (Huang et al., 2009; Martinez et al., 2015). However, quite how ROS production links mechanistically to the activation of LAP remains a key, unanswered question in the field. In addition, while LAP is dependent on Rubicon and ROS, other non-canonical autophagy processes, such as entosis and drug-induced CASM, are not (De Faveri et al., 2020; Florey et al., 2015; Jacquin et al., 2017), suggesting these molecular players are stimulus-specific inputs, rather than being universally required for CASM. This raises another important, open question regarding a molecular mechanism which might unify the diverse forms of non-canonical autophagy.

In considering a possible universal regulator for non-canonical autophagy, we turned our attention to the V-ATPase as a candidate. V-ATPase is a multi-subunit protein complex that acts as a molecular pump, generating proton gradients within intracellular compartments. It is required to acidify lysosomes, and supports the degradation of autophagosomes and engulfed material. As such, V-ATPase is generally thought to play a terminal, degradative role in autophagy related processes. However, in the context of non-canonical autophagy, we hypothesized that V-ATPase might also play an upstream, activating role, based on several lines of evidence. Firstly, inhibition of V-ATPase by BafA1 blocks CASM induced by LAP, entosis, ionophore and lysosomotropic drug stimulation, TRPML1 activation and STING agonists (Fischer et al., 2020; Florey et al., 2015; Goodwin et al., 2021; Jacquin et al., 2017), indicating a broad but undefined role in non-canonical autophagy. Secondly, a direct interaction has recently been identified between V-ATPase and ATG16L1, during *Salmonella* infection, which can be inhibited by the effector protein SopF (Xu et al., 2019). While this interaction has been defined in the context of xenophagy, it is striking that it depends on the ATG16L1 WD40 CTD, a domain specifically required for non-canonical autophagy, thus hinting at a possible connection to CASM. Finally, and in line with this reasoning, STING activation induces non-canonical autophagy in a SopF-sensitive fashion, in a process thus termed V-ATPase–ATG16L1 induced LC3B lipidation (VAIL) (Fischer et al., 2020). Integrating these diverse observations, we hypothesised that V-ATPase may represent a universal regulator of non-canonical autophagy.

In this study, we investigate the role of V-ATPase in CASM, identifying a requirement for enhanced V0-V1 association and direct ATG16L1 engagement across a wide range of non-canonical autophagy processes. We also define the interrelationships between V-ATPase, NADPH Oxidase and ROS during LAP, providing new mechanistic insight into this specific process. Finally, we explore the roles of V-ATPase and non-canonical autophagy during *Salmonella* infection, concluding that CASM represents a host pathogen response, evaded by the SopF effector.

## Results

### V-ATPase is recruited to phagosomes prior to LC3 lipidation

During LAP, the fusion of phagosomes with late endosomes/lysosomes, and their associated acidification via V-ATPase, is assumed to occur after the recruitment of LC3, as a terminal step (Martinez et al., 2015). However, inhibition of V-ATPase with Bafilomycin A1 (BafA1) blocks LC3 recruitment during LAP, and other non-canonical autophagy processes, suggesting V-ATPase may play an additional, upstream role (Florey et al., 2015). To investigate this notion, we revisited the dynamics of LAP using live imaging.

First, fluorescently tagged markers of late endosomes/lysosomes were monitored, in relation to GFP-hLC3A, during phagocytosis of opsonised zymosan particles (LAP). Strikingly, we observed early recruitment of the late endosome marker, RFP-Rab7, to newly formed phagosomes, occurring prior to GFP-hLC3A recruitment (Fig. 1 a; and Video 1). Similarly, the lysosome marker, LAMP1-RFP, is also recruited to phagosomes before GFP-hLC3A (Fig. 1 b; and Video 2). By measuring the duration between onset of markers, we confirmed that phagosomes mature through a Rab7 – LAMP1 – LC3 sequence (Fig. 1 c). The same maturation profile was also observed during entosis, another macro-scale engulfment process that utilises non-canonical autophagy (Fig. S1). Together, these data imply that late endosomes/lysosomes fuse with the phagosome prior to the initiation of non-canonical autophagy signalling.

**Figure 1.**
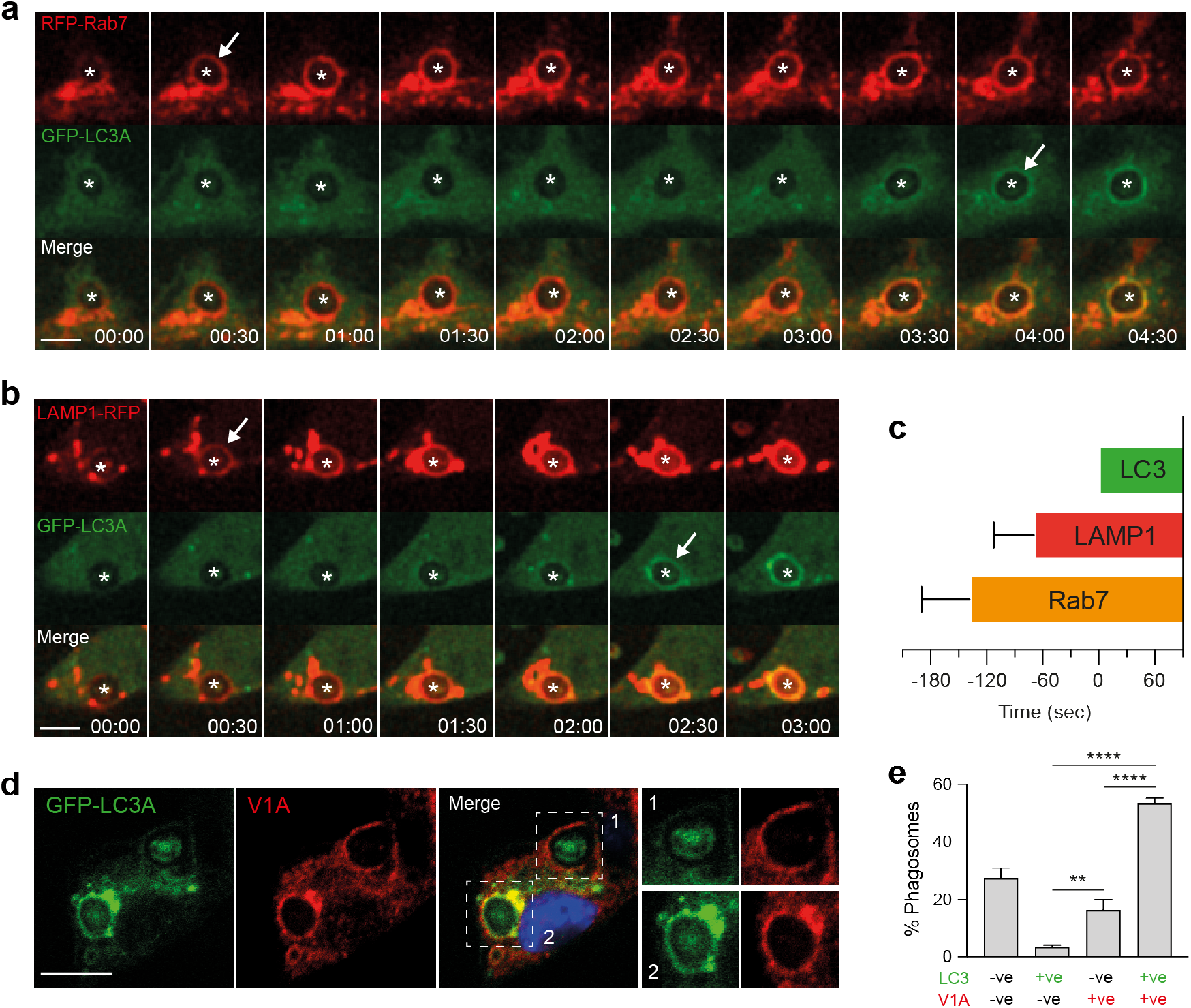
Rab7, LAMP1 and V-ATPase recruit prior to LC3 during LAP. **(a and b)** Representative timelapse confocal microscopy images of opsonised zymosan (OPZ) phagocytosis in RAW264.7 cells expressing GFP-hLC3A and (a) RFP-Rab7 or (b) LAMP1-RFP. Asterisks denotes phagosomes and arrows mark acquisition of fluorescent markers. Scale bar, 5 μm, min:sec. **(c)** Quantification of fluorescent marker acquisition from 10 phagosomes in relation to GFP-hLC3A (time 0). Data represent mean +/- SD. **(d)** Confocal images of OPZ containing phagosomes stained for GFP-hLC3 and ATP6V1A. Scale bar, 5 μm. **(e)** Quantification of phagosome markers from (d). Data represent mean +/- SEM from 3 independent experiments with >100 phagosomes analysed per experiment. ****p<0.0001, **p<0.001, one-way ANOVA followed by Tukey multiple comparison test.

We next sought to visualise V-ATPase directly. This large, multi-subunit protein complex can be challenging to monitor in live cells, as fluorescent tags may interfere with function. Therefore, to assess V-ATPase and LC3 recruitment to phagosomes, immunofluorescent staining was performed at specific timepoints on fixed cells. At 20 minutes post zymosan incubation, the majority of phagosomes were positive for both the V1A subunit of V-ATPase and GFP-hLC3A (Fig. 1 d and e). Importantly, significantly more V1A positive, GFP-hLC3A negative phagosomes were detected than V1A negative, GFP-hLC3A positive (Fig. 1d and e), suggesting that V1A is likely to be recruited first. We can exclude the possibility that GFP-hLC3A has transiently translocated before fixation, as GFP-hLC3A clearly persisted on phagosomes for longer than 20-minutes in live imaging studies. Therefore, our data indicate that, like Rab7 and LAMP1, V-ATPase is present on phagosomes prior to LC3. This is consistent with previous work indicating that V-ATPase recruits to phagosomes at an early stage of their maturation (Sun-Wada et al., 2009).

Together, these data indicate that components of the late endosome/lysosome network, including V-ATPase, are recruited to phagosomes before the onset of LC3 lipidation during LAP. These data question the view that V-ATPase performs an exclusively terminal role, and are consistent with a possible upstream, regulatory function.

### NADPH oxidase and V-ATPase activities are both required for LAP

A major molecular mechanism implicated in LAP involves NADPH oxidase-mediated ROS generation (Martinez et al., 2015); however, the specific mechanistic role of ROS remains unclear. We next considered the possibility of a relationship between NADPH oxidase and V-ATPase during LAP. Using the well characterised pharmacological inhibitors Diphenyleneiodonium (DPI) and Bafilomycin A1 (BafA1), to inhibit NADPH oxidase or V-ATPase respectively, a reduction in GFP-hLC3A recruitment to phagosomes was detected with both (Fig. 2 a). Notably, analysis of fluorescent intensity indicates that BafA1 yields the more pronounced inhibition (Fig. 2 b). These data confirm that both NADPH oxidase and V-ATPase are required for LAP.

**Figure 2.**
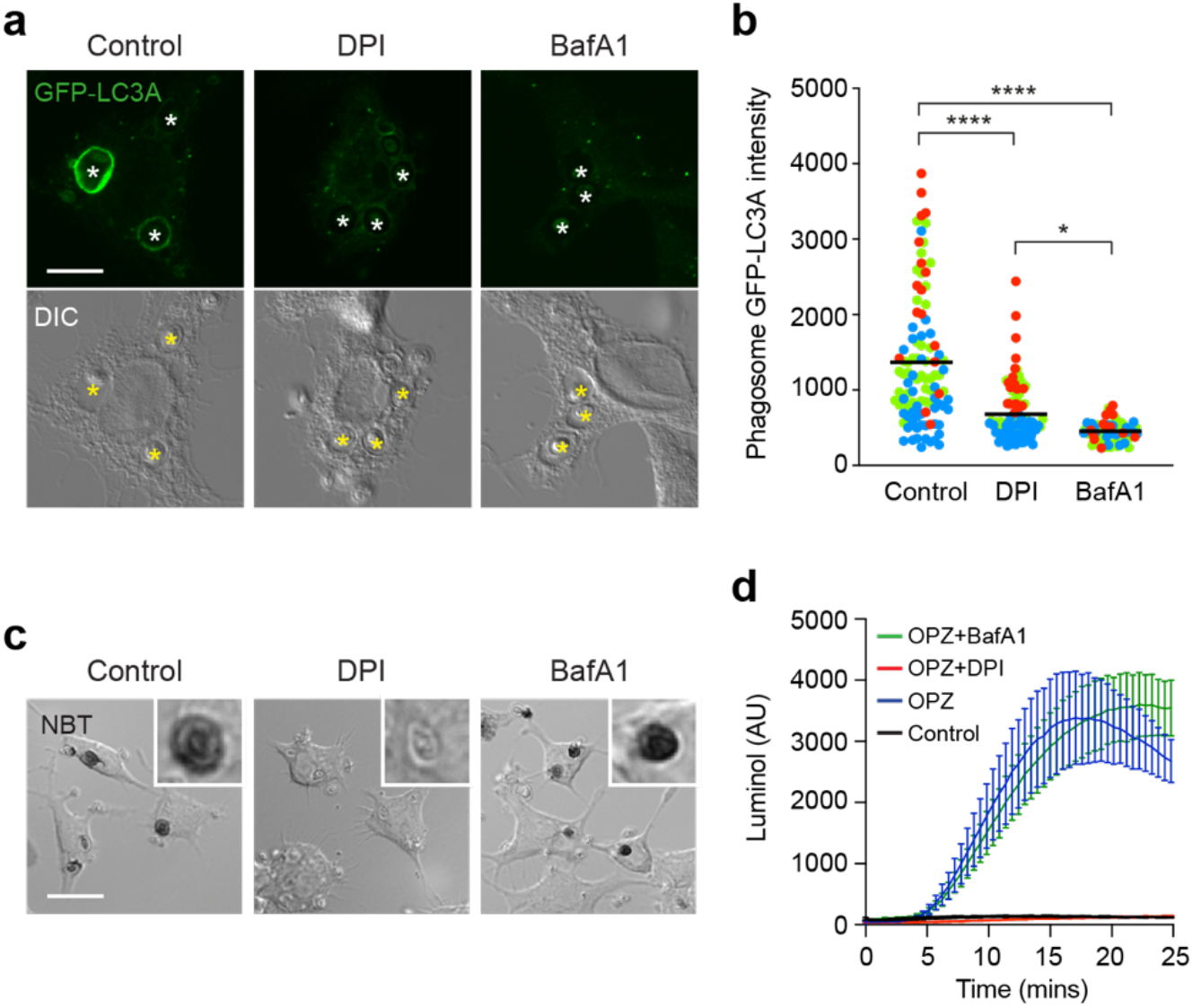
NADPH oxidase and V-ATPase are both required for LAP. **(a)** Confocal DIC and GFP-hLC3A images of OPZ phagocytosis in RAW264.7 cells treated with DPI (5 μM) or BafA1 (100 nM). Asterisk denote phagosomes. Scale bar, 5 μm. **(b)** Quantification of GFP-hLC3A intensity at phagosomes in cells treated as above. Data represent the mean of individual phagosome measurements from 3 independent experiments (red, green, blue); ****p<0.0001, *p<0.03, one-way ANOVA followed by Tukey multiple comparison test. **(c)** Representative confocal DIC images of NBT/formazan deposits in phagosomes from control, DPI and BafA1 treated RAW264.7 cells. Inserts are zoomed in phagosomes. Scale bar, 20 μm. **(d)** Luminol measurements of ROS during OPZ phagocytosis in RAW264.7 cells treated with DPI or BafA1. Data represent the mean +/-SEM from 3 independent experiments.

Next, the effect of each inhibitor on phagosomal ROS generation was analysed, using Nitroblue Tetrazolium (NBT) (Fig. 2 c) and luminol based assays (Fig. 2 d). As shown in Fig. 2 c, O_2_^-^ mediated insoluble formazan deposition, in zymosan containing phagosomes, is abolished by DPI, but not by BafA1 treatment. In agreement with this, DPI treatment blocks zymosan induced ROS production, while BafA1 has no effect (Fig. 2 d). Together, these data establish that while NADPH oxidase is required for ROS generation in phagosomes, V-ATPase activity is not. Moreover, these results show that V-ATPase inhibition blocks LAP, even in the presence of a ROS burst. As such, we can deduce that NADPH oxidase-mediated ROS generation is not sufficient to induce LAP, and that V-ATPase acts either downstream, or in parallel, to promote LC3 lipidation to the phagosome membrane.

### ROS modulates phagosome acidification to drive LAP

To test whether NADPH oxidase might function directly upstream of V-ATPase during LAP, the relationship between ROS and phagosome acidification was assessed, using Lysotracker and live cell confocal microscopy, in the presence or absence of DPI. Strikingly, inhibition of NADPH oxidase significantly increases Lysotracker intensity, signifying increased acidification of zymosan containing phagosomes by V-ATPase (Fig. 3 a). These data agree with previous studies (Mantegazza et al., 2008; Savina et al., 2006), which proposed that production of phagosomal ROS can constrain acidification, by increasing membrane permeability or through the consumption of protons (H+) (Westman and Grinstein, 2020). According to this model, in the absence of ROS, more protons can accumulate, and thus the phagosome is more readily acidified.

**Figure 3.**
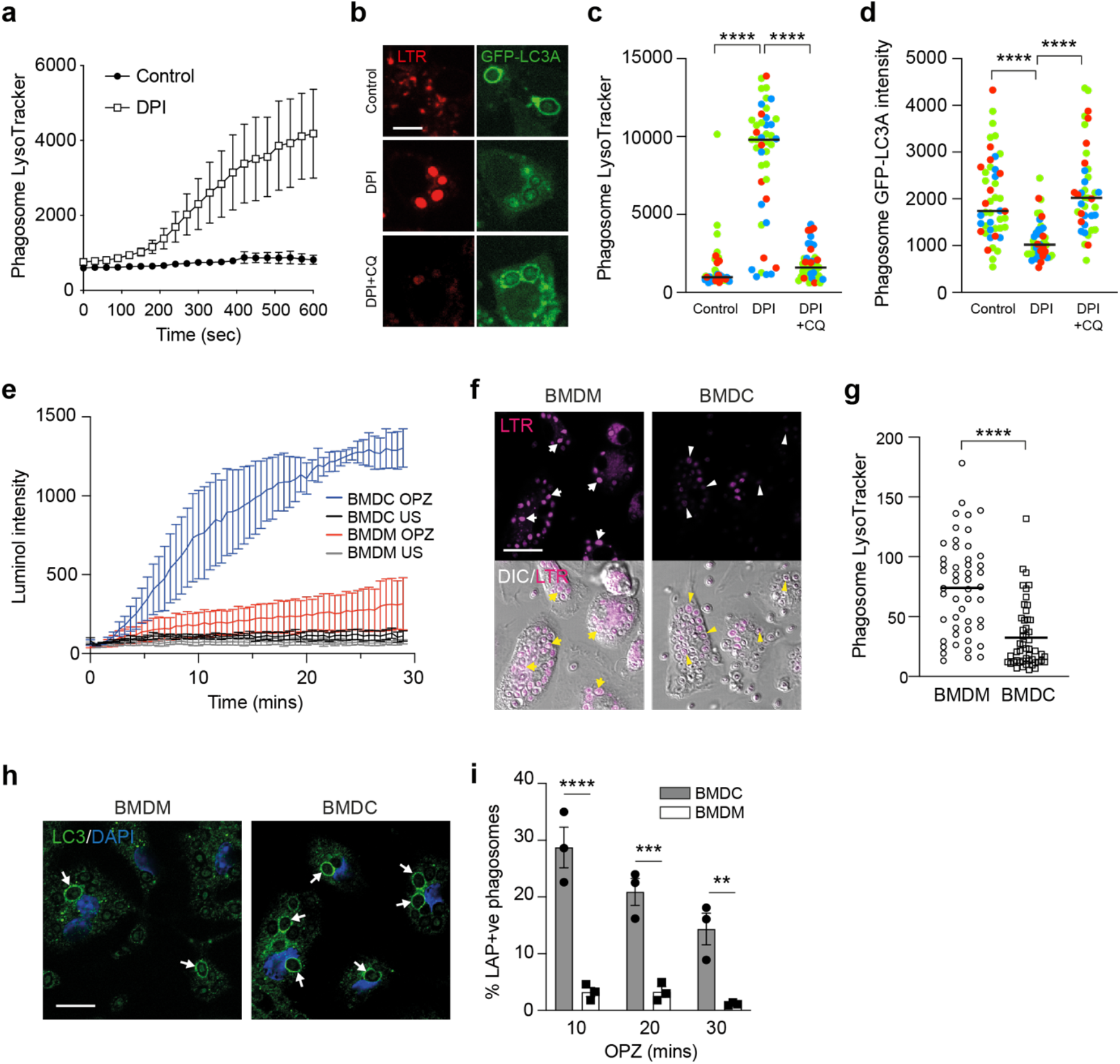
LAP induction involves elevated ROS and raised pH in phagosomes. **(a)** Confocal microscopy measurement of phagosome LysoTracker intensity over time in RAW264.7 cells stimulated with OPZ +/- DPI (5 μM) pre-treatment. Data represent mean +/-SEM of 9 individual phagosomes across multiple independent experiments. **(b)** RAW264.7 cells were pre-treated with DPI (5 μM) or DPI + CQ (100 μM) prior to stimulation with OPZ for 25 min. Representative confocal images of LysoTracker and GFP-hLC3A are shown. Scale bar, 5 μm. **(c and d)** Quantification of (c) Lysotracker and (d) GFP-hLC3A at phagosomes in cells treated as in (b). Data represent the mean of individual phagosome measurements from 3 independent experiments (red, green, blue). ****p<0.0001, one-way ANOVA followed by Tukey multiple comparison test. **(e)** Luminol measurements of ROS during OPZ phagocytosis in primary murine BMDC and BMDM cells. Data represent the mean +/-SEM from 3 independent experiments. **(f)** Representative confocal images of LysoTracker staining and DIC in BMDC and BMDM cells after stimulation with OPZ for 25 min. Arrows denote phagosomes in BMDMs, arrowheads denote phagosomes in BMDCs. Scale bar, 20 μm. **(g)** Quantification of phagosome Lysotracker intensity in cells treated as in (f). Data represent the mean of 50 individual phagosomes. ****p<0.0001, unpaired t test. **(h)** Representative confocal images of BMDC and BMDM cells stimulated with OPZ for 25 min and stained for LC3. Arrows denote LC3 positive phagosomes. Scale bar, 10 μm. **(i)** Quantification of LC3 positive phagosomes from cells stimulated with OPZ for the indicated times. Data represent mean +/-SEM from 3 independent experiments. ***p<0.0001, ***p<0.0003, **p<0.003, two-way ANOVA with multiple comparisons.

To explore this further, we utilised chloroquine (CQ), a weak-base amine, which becomes protonated and accumulates in acidifying compartments, thereby ‘trapping’ H+ ions. We reasoned that CQ could artificially sequester H+, in DPI-treated cells, providing an alternative way to limit protons, in the absence of ROS. In line with our predictions, the addition of CQ countered the DPI-mediated increase in Lysotracker staining (Fig. 3 b and c). Importantly, CQ treatment also rescued GFP-hLC3A recruitment in the presence of DPI, connecting ROS generation and proton concentration directly to LAP (Fig. 3 b and d). Together, these data establish a relationship whereby high phagosomal ROS levels raise pH and drive LAP.

To interrogate this relationship in the absence of pharmacological manipulation, bone marrow-derived murine macrophage (BMDM) and dendritic cells (BMDC) were compared. Previous studies established that mouse BMDC phagosomes generate higher ROS levels, and have a more neutral pH, than BMDM phagosomes (Mantegazza et al., 2008; Savina et al., 2006). Using the luminol assay to measure ROS, and Lysotracker staining to monitor acidification, we confirmed these observations (Fig. 3 e-g). Strikingly, we also found that BMDCs support higher levels of LAP than BMDMs (Fig. 3 h and i), consistent with our model.

Thus, using both pharmacological and cell type-based manipulations, our data indicate that increased LAP is associated with higher ROS production and pH. These findings provide new insight into the molecular mechanisms underlying LAP and uncover a clear link between NADPH oxidase and V-ATPase activities.

### ROS modulates V-ATPase subunit recruitment

We next sought to understand how ROS alters V-ATPase function, and how this results in the lipidation of LC3 to phagosomes. While modulation of phagosome pH does occur with ROS, our data do not support this as a direct mechanism. Firstly, while DPI and BafA1 both inhibit LAP, they yield different effects on phagosomal pH (Fig. S2). Secondly, while BafA1 blocks LAP, monensin and many other drugs that raise lysosomal pH, instead activate non-canonical autophagy (Jacquin et al., 2017). We therefore considered whether other, pump-independent features of V-ATPase might connect ROS and LAP, focussing first on V-ATPase subunit localisation and recruitment.

During LAP, we observe robust recruitment of ATP6V1A, a subunit of the V-ATPase V1 sector, to zymosan containing phagosomes (Fig. 4 a). Notably, both DPI and BafA1 treatment reduce this ATP6V1A signal intensity (Fig. 4 a and b). This result may at first appear somewhat counter intuitive, given that DPI increases the acidification of phagosomes (Fig. 3 a). However, we speculate that in the absence of H+ consumption by ROS, the phagosome requires less V-ATPase activity to support acidification, and therefore recruits less ATP6V1A. In support of this interpretation, artificial sequestration of H+ by CQ reverses the DPI induced reduction in ATP6V1A staining (Fig. 4 c). To extend these observations, the recruitment of ATP6V1A to phagosomes was also compared in BMDCs and BMDMs. Consistent with the above data, BMDC phagosomes exhibit greater ATP6V1A recruitment than BMDMs (Fig. 4 d and e).

**Figure 4.**
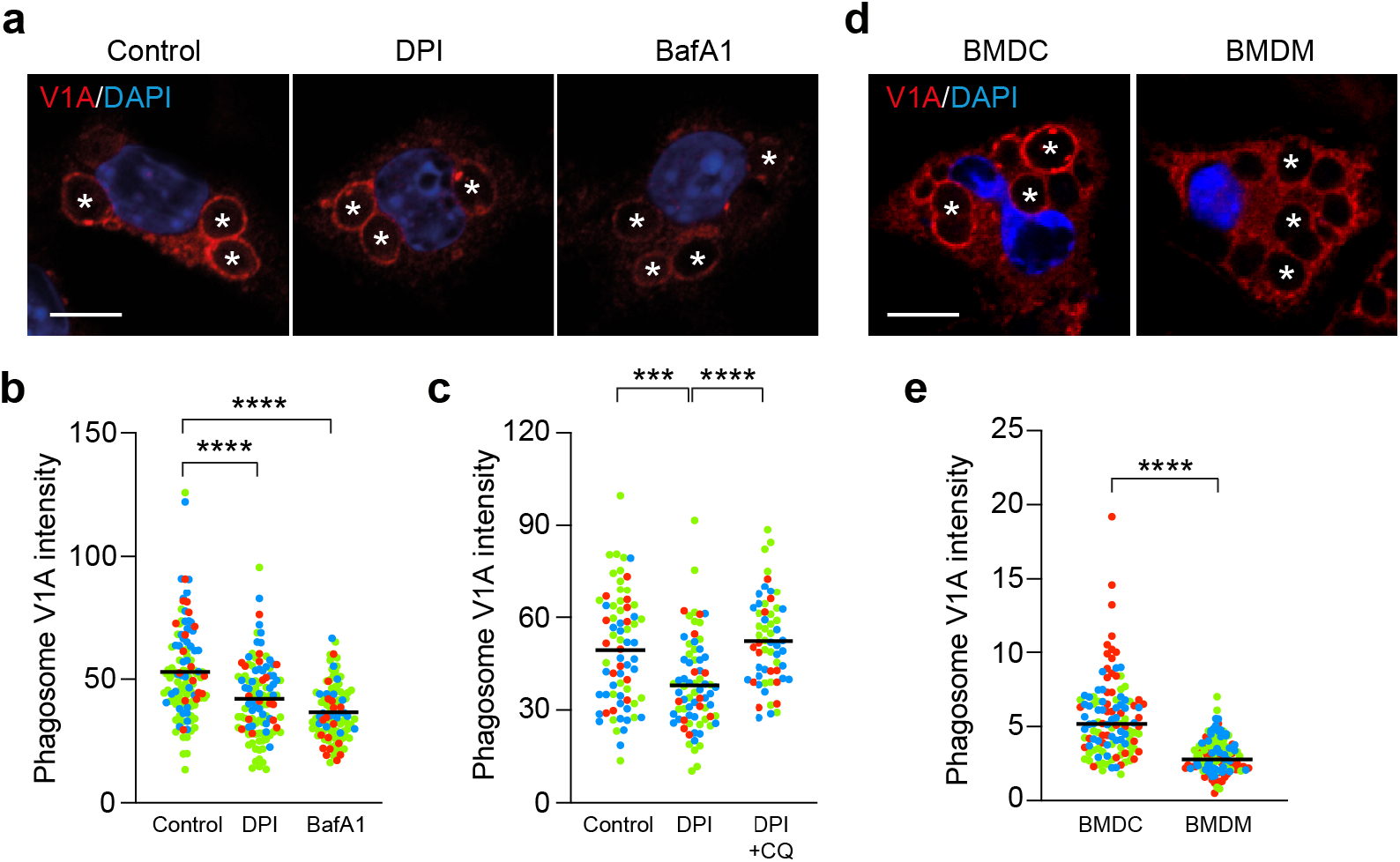
Modulation of ROS regulates phagosomal V-ATPase recruitment during LAP. **(a)** Representative confocal images of ATP6V1A following 25 min OPZ stimulation of RAW264.7 cells pre-treated with either DPI (5 μM) or BafA1 (100 nM). Asterisks denote phagosomes. Scale bar, 5 μm. **(b)** Quantification of ATP6V1A phagosome intensity from cells treated as in (a). Data represent the mean of individual phagosome measurements from 3 independent experiments (red, green, blue). ****p<0.0001, one-way ANOVA followed by Tukey multiple comparison test. **(c)** Quantification of ATP6V1A phagosome intensity from RAW264.7 cells pre-treated with DPI or DPI + CQ. Data represent the mean of individual phagosome measurements from 3 independent experiments (red, green, blue). ****p<0.0001, one-way ANOVA followed by Tukey multiple comparison test. **(d)** Representative confocal images of ATP6V1A following 25 min OPZ stimulation of BMDC or BMDM cells. Asterisks denote phagosomes. Scale bar, 5 μm. **(e)** Quantification of ATP6V1A phagosome intensity from BMDC and BMDM cells. Data represent the mean of individual phagosome measurements from 3 independent experiments (red, green, blue). ****p<0.0001, one-way ANOVA followed by Tukey multiple comparison test.

Taken together, these findings indicate that LAP activation correlates with V-ATPase V1 sector recruitment to the phagosome, which can be regulated by NADPH oxidase activity.

### Increased V0-V1 association drives non-canonical autophagy

V-ATPase activity can be regulated by the association of the cytosolic V1 sector with the membrane bound V0 sector to form a functional holoenzyme, a requisite for pump activity (Collins and Forgac, 2020). To address whether regulated V0-V1 association might play a role in non-canonical autophagy, two structurally distinct V-ATPase inhibitors, BafA1 and Saliphenylhalamide (SaliP), were exploited. These inhibitors bind to distinct sites on V-ATPase, both blocking pump activity (Xie et al., 2004), and raising lysosomal pH. Importantly, however, while BafA1 acts to dissociate V0-V1, SaliP instead drives their association through a covalent adduct formation (Fig. 5 a) (Kissing et al., 2015; Xie et al., 2004). As such, we can use these inhibitors to differentially manipulate V0-V1 association.

**Figure 5.**
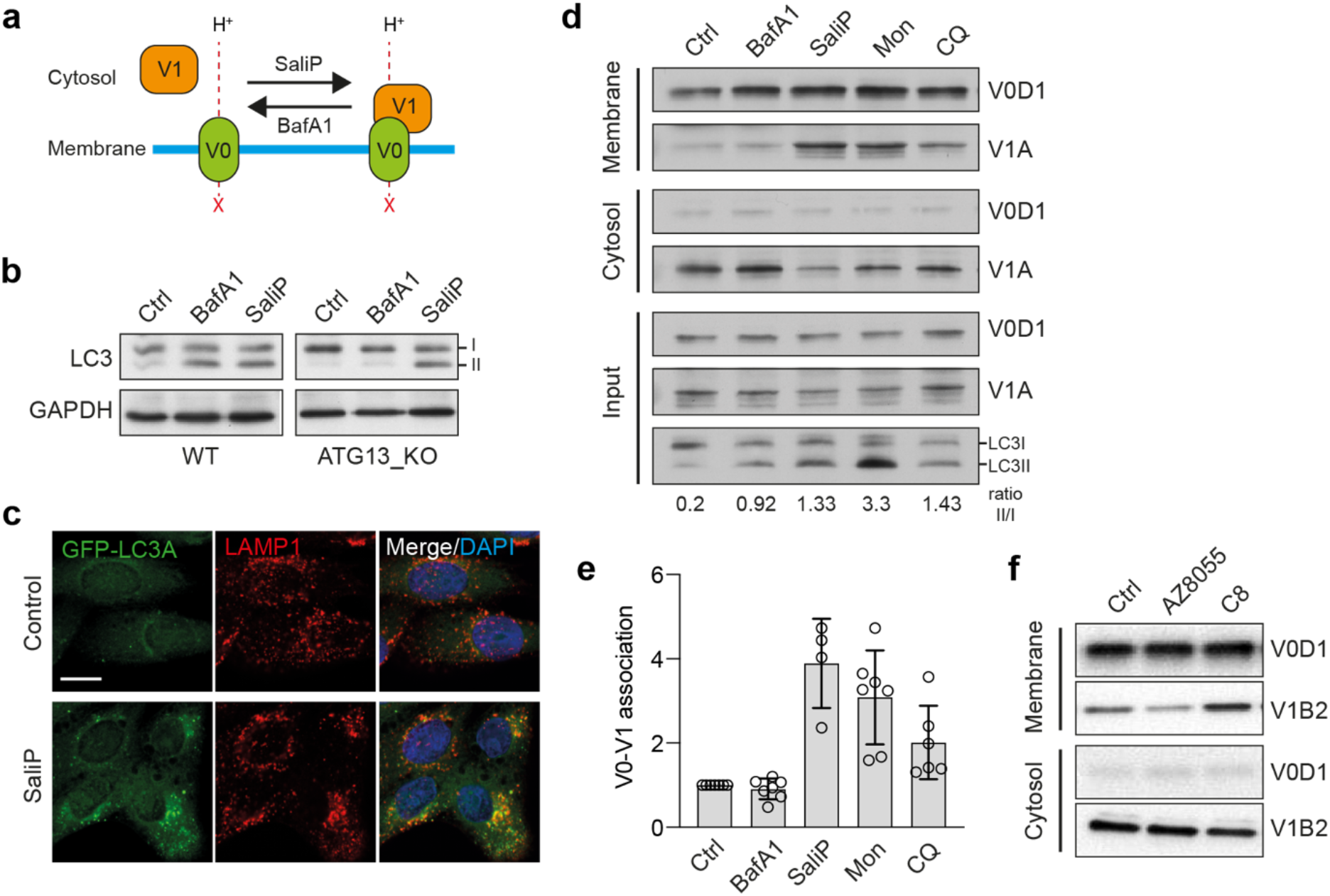
V-ATPase V0-V1 engagement is associated with, and sufficient for, non-canonical autophagy activation. **(a)** Cartoon showing the differential effects of BafA1 and SaliP on V0-V1 association. **(b)** Wild type and *ATG13*^-/-^ MCF10A cells were treated with BafA1 (100 nM) or SaliP (2.5 μM) for 1 h. Western blotting was performed to probe for LC3 (I and II forms marked) and GAPDH. **(c)** Representative confocal images of *ATG13*^-/-^ MCF10A cells stained for LAMP1 and GFP-hLC3A following treatment with SaliP (2.5 μM) for 1 h. **(d)** Wild type MCF10A cells were treated with BafA1 (100 nM), SaliP (2.5 μM), monensin (100 μM) or CQ (100 μM) for 1 h. Following fractionation, input, membrane and cytosol fractions were probed for ATP6V1A, ATP6V0D1 and LC3 by western blotting. LC3II/LC3I ratios shown below. **(e)** Quantification of V0-V1 association from experiments as shown in (d). Data represent mean +/-SD from 4-7 independent experiments. **(f)** HeLa cells were stimulated with either mTor inhibitor AZD8055 (1 μM) or TRPML1 agonist C8 (2 μM) for 90 min. Following fractionation, membrane and cytosol fractions were probed for ATP6V1B2, ATP6V0D1 by western blotting.

Treatment of wild type MCF10A cells with BafA1 or SaliP leads to an increase in GFP-hLC3A lipidation (Fig. 5 b), which would typically be attributed to the inhibition of autophagic flux (Yamamoto et al., 1998). Consistent with this, in ATG13KO cells, which are deficient in canonical autophagy, BafA1 yields no such increase in GFP-hLC3A lipidation. Strikingly, however, SaliP still induces robust GFP-hLC3A lipidation, co-localised with LAMP1, in the autophagosome-deficient ATG13 KO cells (Fig. 5 b and c). These data indicate that SaliP induces GFP-hLC3A lipidation through CASM, and that forced V0-V1 association may be sufficient to trigger non-canonical autophagy.

To investigate this notion further, V0-V1 association was monitored by membrane fractionation in response to known inducers of CASM. Strikingly, both monensin and CQ, two distinct pharmacological agents which disrupt lysosomal ion balance to activate non-canonical autophagy, promote increased V0-V1 association (Fig. 5 d and e). Furthermore, a TRPML1 agonist (C8), which also drives CASM (Goodwin et al., 2021), similarly enhances V0-V1 binding (Fig. 5 f). Notably however, activation of canonical autophagy, via mTOR inhibition (AZD8055), does not increase V0-V1 engagement, suggesting this mechanism is specific to the non-canonical pathway.

Taken together, these data indicate that enhanced V0-V1 engagement is a conserved feature of non-canonical, but not canonical, autophagy processes. Based on the use of SaliP, forced association of V0-V1 is sufficient for CASM activation, and independent of V-ATPase pump activity. We conclude that non-canonical autophagy depends instead upon physical subunit engagement, likely associated with structural changes within the V-ATPase complex itself.

### V0-V1 association drives V-ATPase - ATG16L1 interaction

A direct interaction between V-ATPase and ATG16L1 was recently identified in the context of xenophagy (Xu et al., 2019). Intriguingly, this interaction involves the WD40 CTD of ATG16L1, a domain essential for CASM (Fletcher et al., 2018). We thus hypothesized that V-ATPase, and specifically V0-V1 engagement, may drive CASM via a comparable direct recruitment of ATG16L1. To test this, co-immunoprecipitation experiments were performed in cells expressing either FLAG-tagged WT ATG16L1, or a WD40 CTD point mutant (K490A), which is specifically deficient in non-canonical autophagy (Durgan et al., 2021; Fletcher et al., 2018). Under resting conditions, ATG16L1 does not interact with V-ATPase. However, during LAP, we observe a robust interaction between WT ATG16L1 and the V-ATPase ATP6V1A subunit (Fig. 6 a and e). Importantly, this interaction is completely abolished by the K490A mutation, consistent with the molecular dependencies of non-canonical autophagy, while both wild type and K490A ATG16L1 are able to pull down ATG5. This K490-dependent interaction between ATG16L1 and ATP6V1A is similarly induced by diverse non-canonical autophagy activators including the STING ligand DMXAA (Fig. 6 b and e), TRPML1 activation with C8 (Fig. 6 c and f), monensin (Fig. S3) and influenza A virus infection (data in submission). Strikingly, however, activation of canonical autophagy via mTOR inhibition (PP242), does not induce ATG16L1 – V-ATPase interaction (Fig. 6 d and e), suggesting again that this mechanism is associated specifically with CASM. Finally, we find that treatment of cells with SaliP can also promote V-ATPase-ATG16L1 binding, dependent on K490A, indicating that enhanced V0-V1 association is sufficient to drive this interaction (Fig. 6 g).

**Figure 6.**
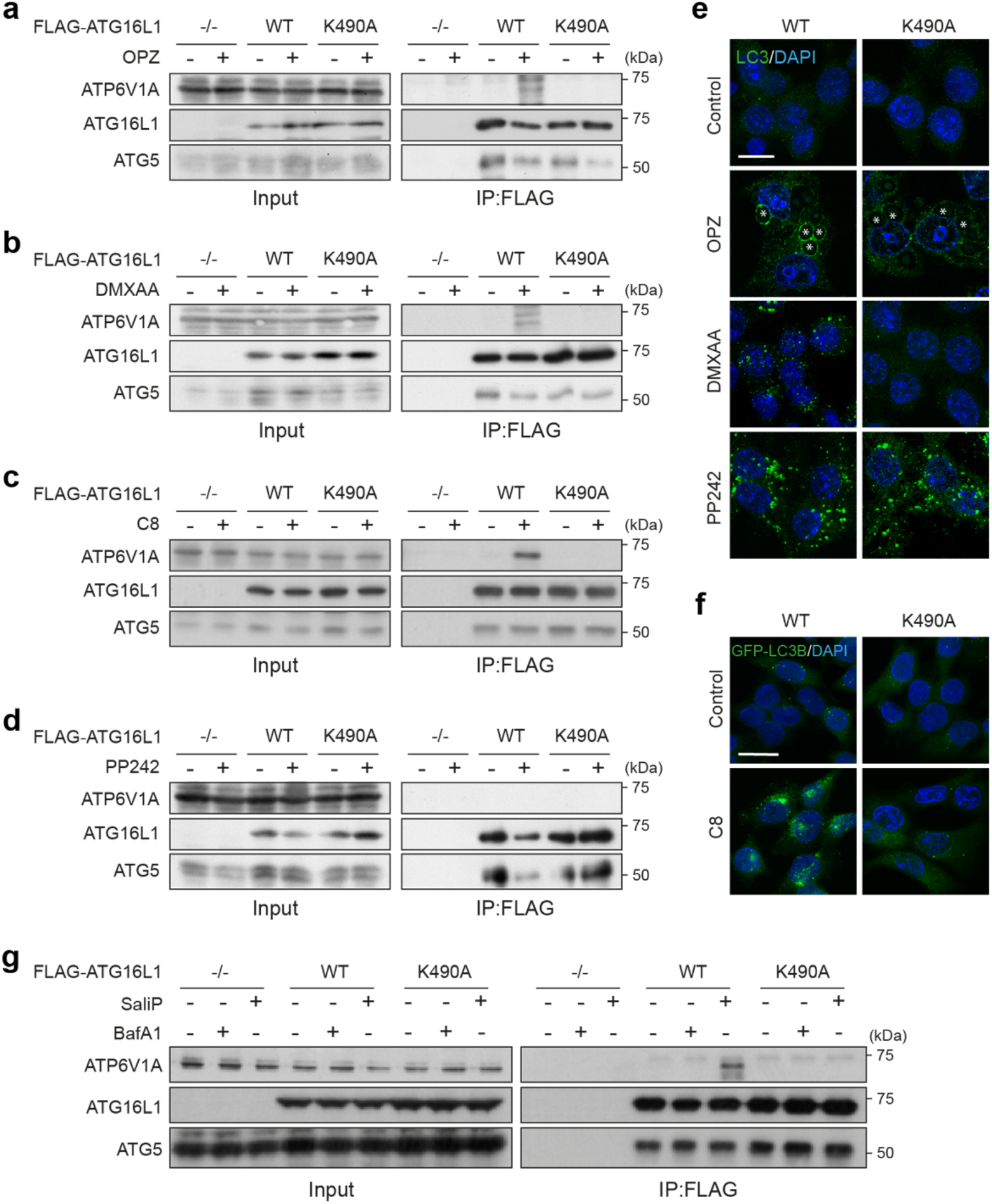
ATG16L1 interacts with V-ATPase during non-canonical autophagy. **(a and b)** *ATG16L1*^-/-^ RAW264.7 cells and those re-expressing Flag-tagged wild type and K490A ATG16L1, were treated with (a) OPZ for 25 min or (b) STING agonist DMXAA (50 μg/ml) for 1 h. Input lysates and Flag immunoprecipitations were probed for ATP6V1A, ATG16L1 and ATG5 by western blotting. (c) *ATG16L1*^-/-^ HCT116 cells and those re-expressing FLAG-tagged wild type and K490A ATG16L1, were treated with TRPML1 agonist C8 (2 μM) for 30 min. Input lysates and FLAG immunoprecipitations were probed by western blotting as above. (d) *ATG16L1*^-/-^ RAW264.7 cells and those re-expressing FLAG-tagged wild type and K490A ATG16L1, were treated with mTor inhibitor PP242 (1 μM) for 1 h. Input lysates and Flag immunoprecipitations were probed by western blotting as above. (e) Confocal images of RAW264.7 cells expressing wild type or K490A ATG16L1 treated with OPZ (25 min), DMXAA (1 h) or PP242 (1 h) and stained for LC3. Asterisks denote phagosomes. Scale bar, 10 μm. (f) Confocal images of GFP-rLC3B HCT116 cells expressing wild type or K490A ATG16L1 treated with C8 (30 min). Scale bar, 15 μm. (g) *ATG16L1*^-/-^ HCT116 cells and those re-expressing FLAG-tagged wild type and K490A ATG16L1, were treated with BafA1 (100 nM) or SaliP (2.5 μM) for 1 h. Input lysates and FLAG immunoprecipitations were probed by western blotting as above.

Together, these data reveal that V-ATPase directly recruits ATG16L1 during diverse non-canonical autophagy processes to drive CASM. We speculate that V0-V1 association permits this interaction, perhaps via a conformational change, and that it requires that the K490 containing pocket, on the top face of the ATG16L1 C terminal WD40 β barrel. We propose that during CASM, the V-ATPase complex may play an analogous role to WIPI2 in canonical autophagy, recruiting ATG16L1 to the appropriate membrane, to specify the site of ATG8 conjugation.

### SopF blocks LAP and non-canonical autophagy

SopF is a *Salmonella* effector protein that blocks the interaction between V-ATPase and ATG16L1 during xenophagy, by ribosylation of Gln124 in the ATP6V0C subunit (Xu et al., 2019). We reasoned that if V-ATPase drives LAP through recruitment of ATG16L1, then this would also be sensitive to SopF. Strikingly, transient overexpression of mCherry-SopF in RAW264.7 cells completely inhibits GFP-hLC3A lipidation during phagocytosis of zymosan (Fig. 7 a and b), without affecting ROS production in phagosomes, as determined by NBT test (Fig. 7 c). These data are consistent with a model in which ROS acts upstream of V-ATPase to modulate ATG16L1 recruitment. Interestingly, stable expression of SopF in MCF10A cells does not interfere with V0-V1 association, as promoted by monensin or SaliP (Fig. 7 d), suggesting it specifically blocks downstream ATG16L1 engagement. Notably, SopF also appears to inhibit non-canonical LC3 lipidation in the context of STING or TRPML1 activation and influenza A virus infection (Fischer et al., 2020; Goodwin et al., 2021; Ulferts et al., 2020). Taken together, our data, alongside these recent studies, support the conclusion that V-ATPase is a universal regulator of CASM, which functions to recruit ATG16L1, in a SopF-sensitive fashion.

**Figure 7.**
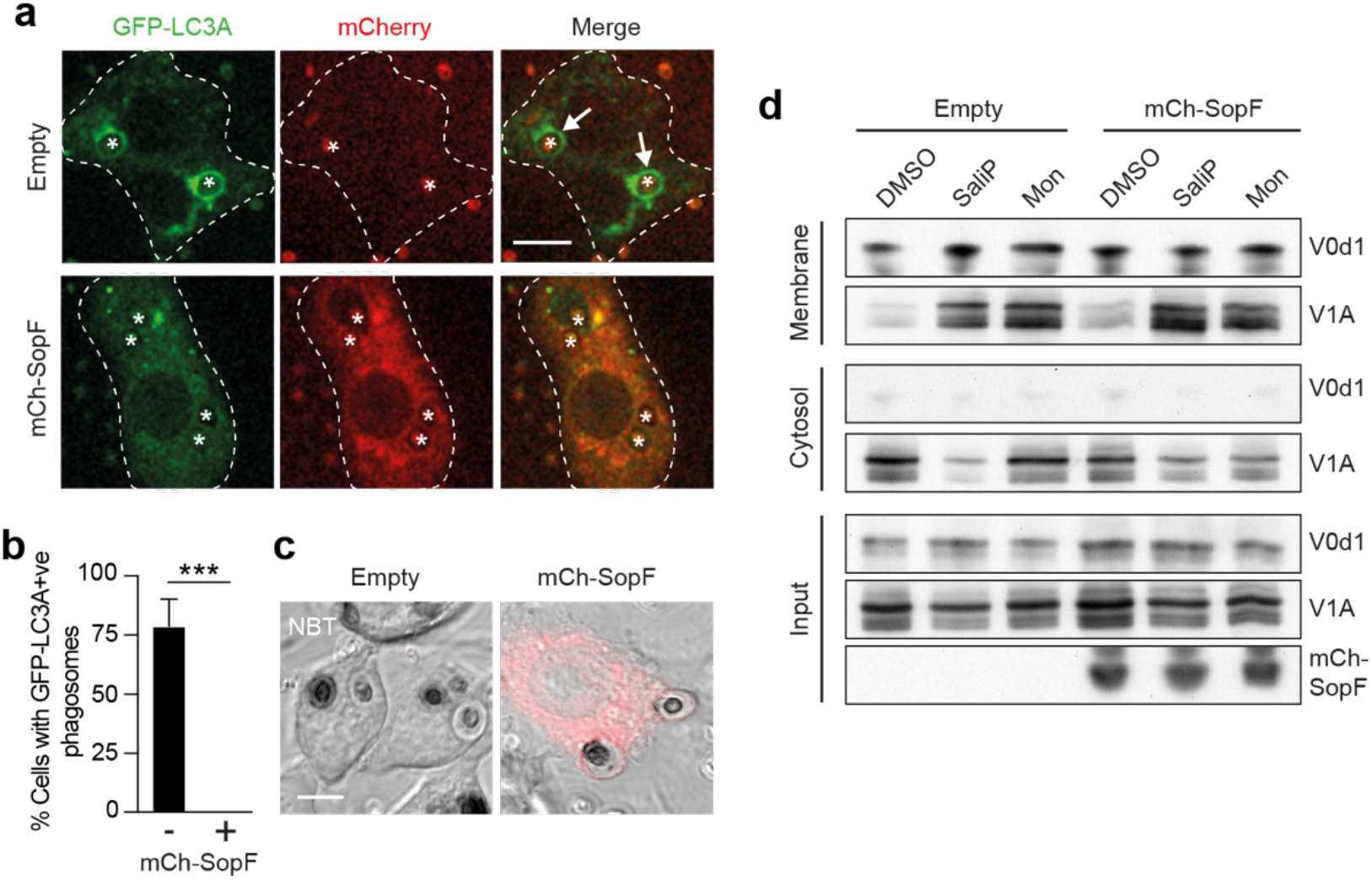
The *Salmonella* effector protein SopF blocks LAP and non-canonical autophagy. **(a)** Confocal images of GFP-hLC3A expressing RAW264.7 cells transfected with mCherry-SopF or empty vector and stimulated with OPZ for 25 min. Asterisks denote phagosomes, arrows mark GFP-LC3A positive phagosomes. Scale bar, 5 μm. **(b)** Quantification of the percentage of phagocytosing mCherry-SopF or empty vector expressing cells that contain GFP-hLC3A positive phagosomes following OPZ stimulation. Data represent the mean +/- SEM from 3 independent experiments. ***p<0.0003, unpaired t test. **(c)** Representative confocal DIC images of NBT/formazan deposits in phagosomes from empty vector and mCherry-SopF expressing RAW264.7 cells. Scale bar, 5 μm. **(d)** Wild type MCF10A cells expressing mCherry-SopF or empty vector were treated with SaliP (2.5 μM) or monensin (100 μM) for 1 h. Following fractionation, input, membrane and cytosol fractions were probed for ATP6V1A, ATP6V0d1 and mCherry by western blotting.

### Non-canonical autophagy contributes to the *Salmonella* host response

Finally, we considered the mechanistic links between V-ATPase, ATG16L1, CASM and SopF in the functional context of host pathogen responses. *Salmonella* SopF provides a mechanism to evade host LC3 lipidation, and this has been elegantly studied in the context of xenophagy (Xu et al., 2019). Here, we investigated the hypothesis that SopF may also suppress CASM as a parallel host response. To explore a possible role for non-canonical autophagy during *Salmonella* infection, HCT116 cells, expressing wild type or K490A ATG16L1, were infected with either wild type or *DsopF Salmonella*. In agreement with Xu *et al*, we found GFP-rLC3B recruitment to *Salmonella* is increased using the *DsopF* strain in wild type cells (Fig. 8 a-c). Importantly, we also found that in non-canonical autophagy deficient ATG16L1 K490A cells, GFP-rLC3B recruitment to either strain is dramatically reduced (Fig. 8 a-c). Considering the K490A mutation has no effect on canonical autophagy, or autophagosome formation (Fletcher et al., 2018; Rai et al., 2019) (Fig. 6 e), these data strongly suggest that *Salmonella* infection also induces a non-canonical autophagy response, which is disrupted by SopF targeting of the V-ATPase-ATG16L1 axis. Together, these findings indicate that *Salmonella* infection triggers CASM as a defensive host response, and that SopF has evolved as a mechanism of evasion.

**Figure 8.**
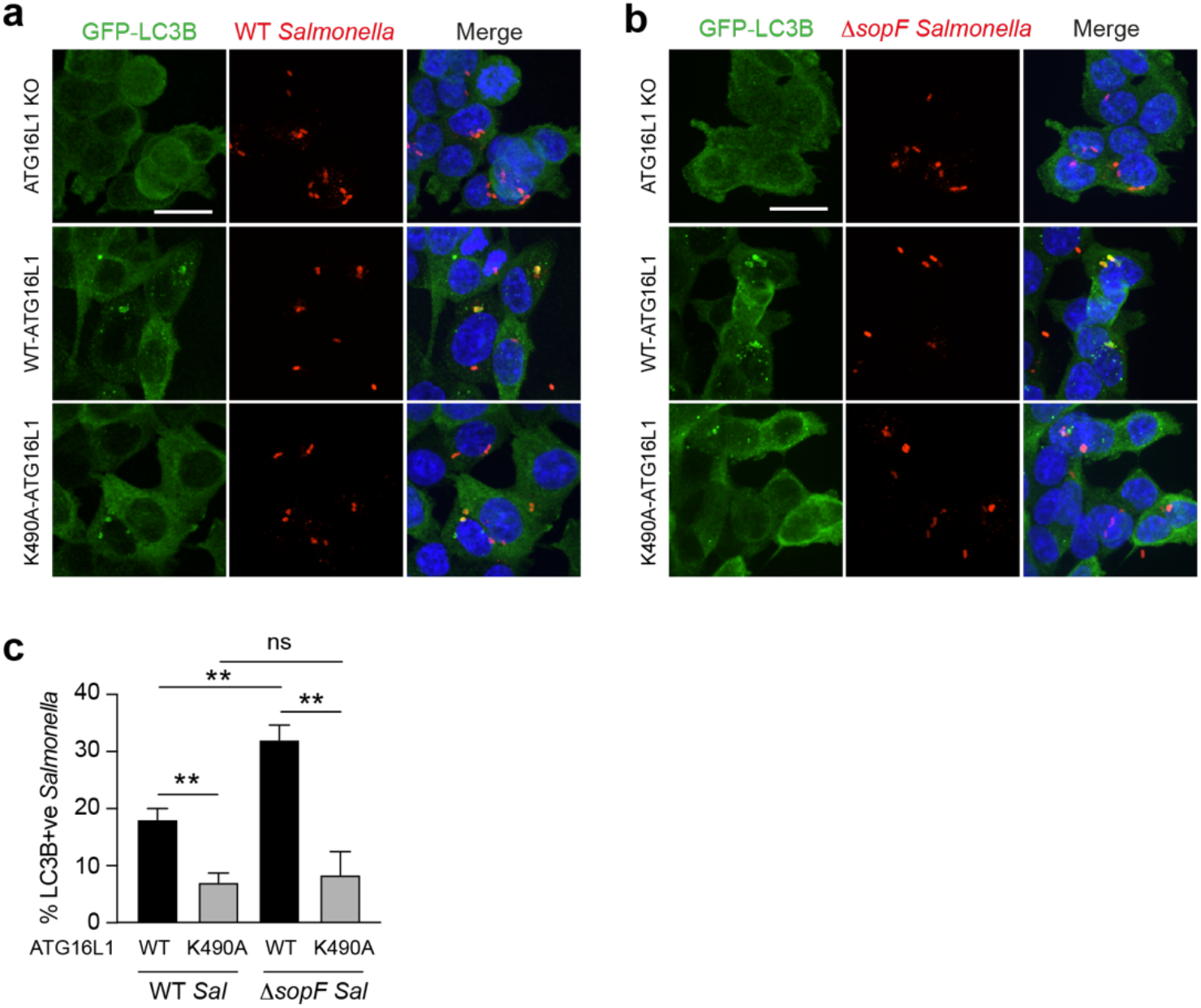
Non-canonical autophagy contributes to the *Salmonella* host response. **(a and b)** *ATG16L1*^-/-^ HCT116 cells and those re-expressing FLAG-tagged wild type and K490A ATG16L1, were infected with either (a) wild type or (b) *ΔsopF Salmonella* for 1 h and imaged by confocal microscopy. Scale bar, 10 μm. **(c)** Quantification of GFP-rLC3B positive *Salmonella* in cells treated as in (a and b). At least 100 bacteria were counted for each condition. Data represent mean +/-SD from 3 independent experiments. **p<0.01, ****p<0.0001, one-way ANOVA followed by Tukey multiple comparison test.

## Discussion

Non-canonical autophagy, or CASM, has emerged as a pathway with vital functions in a wide range of biological processes, including immunity, vision and cancer. A number of molecular players have been implicated in regulation of this pathway, including the core ATG8 conjugation machinery, NADPH oxidase, Rubicon and V-ATPase. However, exactly how these components work together, and whether they are context-specific or common to all forms of CASM, has remained unclear. In this study, we focussed on the involvement of the V-ATPase complex, its interplay with ATG16L1 during non-canonical autophagy processes, and its specific connection to NADPH oxidase/ROS during LAP.

V-ATPase was first implicated in non-canonical autophagy through pharmacological studies that demonstrated opposing effects of BafA1 and CQ on CASM, during LAP and entosis (Florey et al., 2015). BafA1 and CQ are both lysosomal inhibitors, but with different mechanisms of action. BafA1 functions as a V-ATPase inhibitor, and blocks CASM, while CQ, which acts instead to quench protons in the lysosome, induces CASM, in a BafA1 sensitive fashion. These findings indicated that CASM responds to the lumenal ionic balance in lysosomes, and that V-ATPase is essential for the lipidation of ATG8s to single membranes during non-canonical autophagy.

This notion was somewhat unexpected, as V-ATPase is typically thought of as a downstream player during autophagy-related processes, driving terminal lysosomal degradation. However, an upstream, regulatory role for V-ATPase was further reinforced by the finding that, like CQ, a wide range of other ionophores and lysosmotropic agents, including monensin, nigericin, CCCP and clinically used drugs such as lidocaine and amiodarone, similarly induce CASM, in a BafA1 sensitive manner (Jacquin et al., 2017). Furthermore, a comparable pharmacological profile has also been observed during non-canonical autophagy following STING activation, through a process thus termed VAIL (Fischer et al., 2020), and following TRPML1 activation (Goodwin et al., 2021).

Despite the growing evidence that V-ATPase is required to induce non-canonical autophagy, exactly how this complex might sense and respond to different stimuli, and activate CASM, remained unknown. In this study, we have defined a key regulatory mechanism (Fig. 8). Specifically, we find that neutralization of endolysosomal compartments leads to an increased association of the V-ATPase V0-V1 subunits, which in turn facilitates direct ATG16L1 recruitment, through a K490-dependent interaction with the ATG16L1 C terminal WD domain. This mechanism operates in all tested examples of non-canonical autophagy, suggesting this represents a universal regulatory mechanism for the pathway.

Activation of non-canonical autophagy may thus represent a cellular response to stressed or perturbed endolysosomal compartments, that transduce this signal via increased V0-V1 association. It seems likely that these stresses, including lumenal pH change or ionic imbalance, may involve different mechanisms depending on the specific stimulus. For example, pharmacological ionophores and lysosomotropic drugs directly neutralize endolysosomal compartments, which then increase V0-V1 association in an attempt to re-acidify; the ability of V-ATPase to sense and respond to altered lumenal pH in this way has been previously proposed (Marshansky, 2007). In a similar way, the influenza A virus M2 protein, which acts as a H+ proton ionophore, neutralizes intracellular vesicles to activate CASM (Fletcher et al., 2018; Ulferts et al., 2020).

During the activation of LAP, our data indicate that phagosome pH is neutralized by the high levels of ROS generated by NADPH oxidase. The ROS consume protons, thereby raising pH and driving increased phagosomal V1 levels in response. By linking together NADPH Oxidase, ROS and V-ATPase, these findings reveal direct connections between some of the key players in LAP, thus building towards a unifying mechanism. We note that phagosomal ROS have also been linked to the inhibition of ATG4 deconjugation activity during LAP (Ligeon et al., 2021), which may perhaps be related more to prolonging ATG8 lipidation at phagosomes than to its induction.

Further work will be required to assess the triggers for enhanced V0-V1 association in other contexts. In the case of entosis, which involves degradation of an entire internalised cell, it is tempting to speculate that the level of V-ATPase activity required to acidify such a large compartment may necessitate enhanced V0-V1 association. During VAIL, the trafficking of STING to endolysosomes, or TBK1 phosphorylation, may be involved. However, overall, it appears likely that while multiple routes exist to trigger CASM at endolysosomal compartments, these all converge at V-ATPase, and specifically upon enhanced V0-V1 association, as a common step in non-canonical autophagy.

Importantly, a direct interaction between V-ATPase and the WD40 C terminal domain (CTD) of ATG16L1 was recently identified during *Salmonella* xenophagy (Xu et al., 2019). In this study, we report that this interaction is also induced across a range of non-canonical autophagy processes, including LAP, activation of TRPML1 or STING, and drug-induced CASM (monensin), but importantly, not during canonical autophagy. We show that ATG16L1 binding occurs upon enhanced V0-V1 association, in a pump-independent manner, and indeed that increased V0-V1 engagement, triggered by SaliP, is sufficient to recruit ATG16L1. Furthermore, we map the interaction to a specific residue in the ATG16L1 WD40 CTD (K490). Together, these data indicate that the interaction between V-ATPase and ATG16L1 is universally required for non-canonical autophagy. We speculate that V-ATPase may perform a similar function to WIPI2 in canonical autophagy, specifying the site of ATG16L1 recruitment to direct ATG8/LC3 lipidation during CASM.

The *Salmonella* effector SopF ribosylates Gln124 of ATP6V0c in the V-ATPase, inhibiting its interaction with ATG16L1 during xenophagy (Xu et al., 2019), and also during STING activation (Fischer et al., 2020), Influenza virus A infection (Ulferts et al., 2020) and a range of other CASM processes, as shown here. Further to this, our data show that while SopF blocks CASM, it does not block increased V0-V1 association. These findings suggest a specific sequence of events during CASM (Fig. 8), and the existence of a SopF-sensitive conformational change, that occurs during increased V-ATPase activity and V0-V1 association, to engage ATG16L1. Using cryo-electron microscopy, recent advances have been made in determining the complete mammalian structure of V-ATPase (Wang et al., 2020a; Wang et al., 2020b), where different holoenzyme states have been identified. In the future it may be possible to resolve specific structural features in SopF-modified versus unmodified complexes, that are required to support ATG16L1 WD40 CTD binding.

V-ATPase activity is well known to be regulated along the endocytic pathway, with increased assembly and lower pH as it progresses towards lysosomes (Lafourcade et al., 2008). However, the molecular mechanisms regulating V0-V1 association in mammalian cells remain poorly understood. Glucose and amino acid availability can modulate the assembly and association of the V-ATPase complex (McGuire and Forgac, 2018; Stransky and Forgac, 2015), in a PI3-kinase dependent manner, and lipid composition is proposed to play a regulatory role, with PI(3,5)P2 stabilising the V0-V1 association (Li et al., 2014). Changes in V0-V1 levels have also been reported in different differentiation states of murine dendritic cells, with more mature cells displaying increased V0-V1 association (Liberman et al., 2014); our data now reveal an increased V1A recruitment to phagosomes in bone marrow-derived dendritic cells compared to macrophage. Thus, it seems likely that differences in non-canonical autophagy activation may be observed between different cell types and species, dependent on basal differences in their regulation of V-ATPase.

While non-canonical autophagy regulates a wide range of fundamental biological processes (Heckmann et al., 2017), the underlying molecular and cellular functions of CASM are still not fully understood. During LAP, ATG8 recruitment has been proposed to modulate phagosome maturation and content degradation. However, while some studies suggest a role for ATG8s in promoting phagosome maturation and lysosome fusion (Gluschko et al., 2018; Henault et al., 2012; Martinez et al., 2015), others suggest instead that ATG8 recruitment delays maturation to stabilise phagosomes (Romao et al., 2013). In another study, no gross defect in phagosome maturation was detected upon inhibition of ATG8 lipidation (Cemma et al., 2016), and, BafA1, which inhibits LAP, does not impair the recruitment of LAMP1 to phagosomes (Kissing et al., 2015). Considering our new data, which show that ATG8/LC3 is recruited to phagosomes after the acquisition of late endosome/lysosome markers, it seems possible that LAP may act to fine tune, rather than initiate, phagosome maturation. For instance, perhaps ATG8/LC3s recruit interacting proteins to modulate the extent of lysosome fusion and/or signalling from phagosomes. Further work will be required to resolve these debates in the literature and to elucidate possible cell type or context specific differences.

Non-canonical autophagy and the ATG16L1-V-ATPase axis facilitate a key innate immune response to pathogen infection. As such, pathogens have evolved evasion strategies, which further underscore the functional importance of CASM. The intracellular pathogen, *Legionella pneumophila*, expresses an effector protein, RavZ, which deconjugates ATG8 from both PE and PS (Durgan et al., 2021), and blocks LAP (Martinez et al., 2015). Cell wall melanin from the fungal pathogen, *Aspergillus fumigatus*, inhibits LAP by interfering with NADPH oxidase and ROS production (Akoumianaki et al., 2016; Kyrmizi et al., 2018). As noted above, the *Salmonella* effector protein, SopF, directly targets the ATG16L1-V-ATPase interaction to permit bacterial growth (Xu et al., 2019); to date, this effect has been attributed solely to xenophagy inhibition. Here, using genetic models to specifically inhibit non-canonical autophagy, we build on these findings to show that SopF also inhibits CASM, and suggest that the ATG8 response to *Salmonella* is, in part, due to non-canonical autophagy. These data are consistent with other studies implicating LAP in *Salmonella* infection (Masud et al., 2019). In the future it would be of interest to explore the impacts of other pathogen effectors on CASM, such as VopQ from *Vibrio parahaemolyticus* and SidK from *Legionella*, both of which target V-ATPase.

In conclusion, we have defined a key molecular mechanism unifying the various forms of non-canonical autophagy, whereby enhanced V0-V1 association recruits ATG16L1 to V-ATPase positive, endolysosomal compartments to drive CASM. This mechanism explains the unusual pharmacological profile of CASM processes, and provides new molecular insight into how NADPH oxidase and ROS activate LAP. These findings also build on the recent elucidation of V-ATPase-ATG16L1 binding during *Salmonella* induced xenophagy, indicating that this interaction also occurs during non-canonical autophagy, and indeed that V-ATPase specifies the site of single membrane ATG8 lipidation during CASM. Finally, our data implicate CASM as a parallel pathogen response during Salmonella infection, blocked by the effector protein SopF.

**Figure 9.**
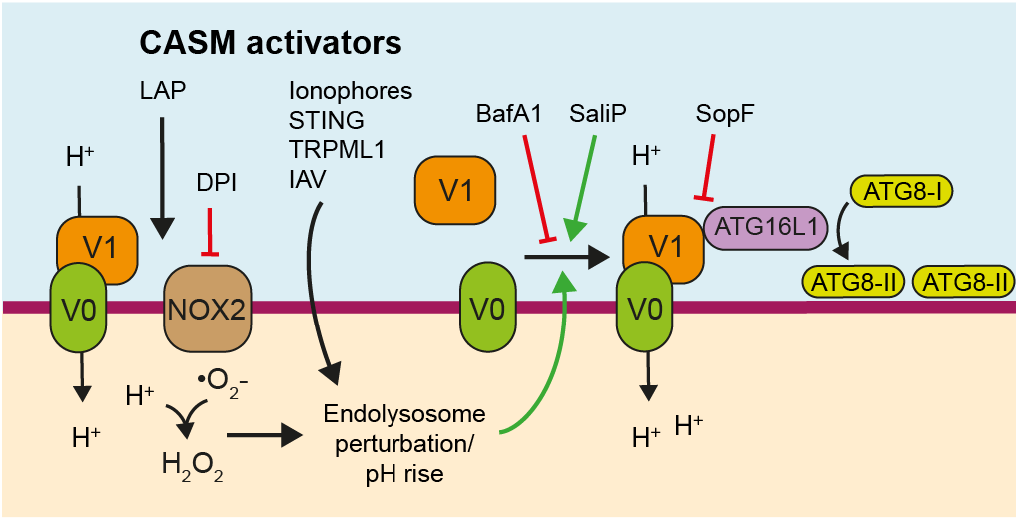
Model of CASM activation. Stimuli that induce perturbations in endolysosomal ion and pH balance drive V0-V1 engagement that promotes the recruitment of ATG16L1 through its WD40 CTD and results in CASM. SopF modifies V-ATPase to inhibit ATG16L1 interaction while BafA1 interferes with V-ATPase activity, both resulting in inhibition of CASM. During LAP, NOX2 dependent ROS production consumes phagosomal H+ protons, in a DPI sensitive manner, which alters phagosome pH and drives the SopF-sensitive interaction between V-ATPase and ATG16L1, which then directs ATG8 lipidation to phagosomes.

## Acknowledgements

We thank Nick Ktistakis, Simon Cook and Phill Hawkins for helpful discussions and critical review of the manuscript, and the BI Imaging facility for all their support. This work was supported by grants from the BBSRC, BB/P013384/1 (BBS/E/B/000C0432 and BBS/E/B/000C0434), BB/R019258/1 and Cancer Research UK Career Development award C47718/A16337. J.H.B. holds the Pitblado Chair in Cell Biology. Work in the lab of J.H.B. was supported by an operating grant from the Canadian Institutes of Health Research (FDN#154329).

## Contributions

K.M.H and E.J. designed and carried out experiments. T.L. and J.H.B. carried out and analysed *Salmonella* experiments. J.M.G provided reagents and insights and performed preliminary experiments. J.D. designed experiments and wrote the manuscript. O.F. designed and carried out experiments and wrote the manuscript.

## Materials and Methods

### Antibodies

Antibodies used were rabbit polyclonal anti-ATG16L1 (8090, Cell Signalling, WB 1:1000), rabbit polyclonal anti-ATG5 (2630, Cell Signalling, WB 1:1000), rabbit polyclonal anti-LC3A/B (4108, Cell Signalling, WB 1:1000, IF 1:100), rabbit monoclonal anti-ATP6V1A (ab199326, Abcam, WB 1:2000, IF 1:250), mouse monoclonal anti-ATP6V0d1 (ab56441, Abcam, WB 1:1000), mouse monoclonal anti-LAMP1 (555798, BD Biosciences, IF 1:100), mouse monoclonal anti-mCherry (ab125096, Abcam, WB 1:1000), mouse monoclonal anti-GAPDH (ab8245, Abcam, WB 1:1000), Alexa Fluor 488 polyclonal goat anti-rabbit IgG (A-11034, ThermoFisher, IF 1:500), Alexa Fluor 568 polyclonal goat anti-mouse IgG (A-11004, ThermoFisher, IF 1:500), Alexa Fluor 568 polyclonal goat anti-rabbit IgG (A-11011, ThermoFisher, IF 1:500), HRP-conjugated anti-rabbit IgG (7074, Cell Signalling, WB 1:2000), HRP-conjugated anti-mouse IgG (7076, Cell Signalling, WB 1:2000).

### Reagents

Reagents and chemicals used were BafA1 (1334, Tocris, 100 nM), PP242 (4257, Tocris, 1 μM), AZD8055 (S1555, Selleckchem, 1 μM), Monensin (M5273, Sigma, 100 μM), DPI (D2926, Sigma, 5 μM), human serum (H2918, Sigma), DMXAA (D5817, Sigma, 50 mg/ml), Zymosan (Z4250, Sigma), NBT (N6876, Sigma), murine IFNg (315-05, Peprotech), DAPI (D9542, Sigma), IN-1 (17392, Caymen, 1 μM), LysoTracker Red DND-99 (L7528, ThermoFisher), LysoTracker Deep Red (L12492, ThermoFisher), anti-FLAG M2 magnetic beads (M8823, Sigma). Saliphenyhalamide (SaliP, 2.5 μM) and TRPML1 agonist C8 (2 μM) were provided by Casma Therapeutics, USA.

### Plasmid constructs

pmCherry-SopF was generated previously (Lau et al., 2019) and kindly provided by Dr Leigh Knodler. mCherry-SopF was cloned into pQCXIP using AgeI/BamHI restriction sites. mRFP-Rab7 was a gift from Ari Helenius (Addgene plasmid # 14436). pBabe-Puro-RFP-LAMP1 was kindly provided by Dr Michael Overholtzer.

### Cell culture

WT or *ATG13*^-/-^ MCF10A cells (human breast epithelial), expressing GFP-LC3A (human), were prepared as described previously (Jacquin et al., 2017) and cultured in DMEM/F12 (Gibco, 11320074) containing 5% horse serum (ThermoFisher, 16050-122), EGF (20 ng/ml; Peprotech AF-100-15), hydrocortisone (0.5 mg/ml; Sigma, H0888), cholera toxin (100 ng/ml; Sigma, C8052), insulin (10 μg/ml; Sigma, I9278), and penicillin/streptomycin (100 U/ml; Gibco 15140-122) at 37°C, 5% CO_2_.

HCT116 cells (human colorectal epithelial) were maintained using DMEM (Gibco, 41966-029) supplemented with 10% FBS (Sigma, F9665) and penicillin/streptomycin (100 U/ml, 100 μg/ml; Gibco 15140-122) at 37°C, 5% CO2. A panel of HCT116 lines expressing GFP-LC3B (rat) and different ATG16L1 constructs, were derived from *ATG16L1^-/-^* cells, reconstituted with pBabe-Puro FLAG-S-*ATG16L1* (wild type or K490A), as described previously (Fletcher et al., 2018).

RAW264.7 cells were obtained from ATCC and maintained in DMEM (Gibco, 41966-029) supplemented with 10% FBS (Sigma, F9665) and penicillin/streptomycin (100 U/ml, 100 μg/ml; Gibco 15140-122) at 37°C, 5% CO_2_. *ATG16L1*^-/-^ were generated and reconstituted with pBabe-Puro FLAG-S-ATG16L1 (wild type or K490A), and GFP-LC3A (human) as described previously (Durgan et al., 2021; Lystad et al., 2019).

### Retrovirus generation and infection

For mCherry-SopF virus infection, MCF10A cells were seeded in a 6 well plate at 5 × 10^4^ per well. The next day 1 ml viral supernatant was added with 10 μg/ml polybrene for 24 h followed by a media change. Cells were then selected with puromycin (2μg/ml).

### BMDC and BMDM isolation

Bone marrow extracted from the hind legs of one C57/BL6 mouse was resuspended in 10 ml complete RPMI 1640 media (Sigma, R8578) (10% FBS, 1% P/S, 50 μM 2-mercaptoethanol) then centrifuged at 1500 rpm for 5 min. The bone marrow was then resuspended in 1 ml of red blood cell lysis buffer (168 mM NH4Cl, 100 μM KHCO3, 10 mM Na2EDTA (pH 8), in milliQ H2O, final pH 7.3) and agitated for 2 min prior to being resuspended in 10 ml complete RPMI 1640 media and centrifuged at 1500 rpm for 5 min. For dendritic cell differentiation the bone marrow was resuspended in 25 ml of complete RPMI 1640 media supplemented with 20 ng/ml GM-CSF (Peprotech #315-03), 10 ng/ml IL-4 (Peprotech #AF-214-14) and 50 ng/ml Amphotericin B (Gibco #15290018). For macrophage differentiation the bone marrow was resuspended in 25 ml of complete RPMI 1640 media supplemented with 20 ng/ml M-CSF (Peprotech #AF-315-02) and 50 ng/ml Amphotericin B. The cells were incubated for 3 days at 37°C, 5% CO2, in low-adherence 90 mm sterilin petri dishes (101R20, ThermoFisher). On the third day media was removed and centrifuged at 1500 rpm for 5 min. The non-adherent cells were then resuspended in fresh differentiation media and added back to the dishes containing the adherent cells for a further 3 days of differentiation. On the sixth day cells non-adherent cells in the media are removed and adherent cells are harvested by gently scraping into complete RPMI 1640 media. The cells were then centrifuged at 1500 rpm for 5 min and seeded in complete RPMI 1640 media without differentiation cytokines.

### Amaxa nucleofection

Transient transfection was performed by electroporation using a Nucleofector II instrument (Lonza, Wakersville, MD, USA) and Lonza nucleofection kit V (Lonza, VCA-1003) following manufacturer’s guidelines. Briefly, 2 x 10^6^ RAW264.7 were electroporated with 2 μg mCherry-SopF or RFP-Rab7 using programme D-032.

### Fluorescent Live cell microscopy and analysis

Prior to imaging, RAW264.7 cells were plated overnight on 35 mm glass-bottomed dishes (MatTek) in the presence of 200 U/ml IFNg. Opzonised zymosan (OPZ) particles were generated by incubating Zymosan A from Saccharomyces (Z4250, Sigma) in human serum (P2918, Sigma) for 30 min at 37°C. The zymosan was then centrifuged at 5,000 rpm for 5 min, and then resuspended in PBS at 10 mg/ml. The solution was passed through a 25-gauge needle using a 1 ml syringe several times to break up aggregates. Cells were mounted on a spinning disk confocal microscope, comprising a Nikon Ti-E stand, Nikon 60x 1.45 NA oil immersion lens, Yokogawa CSU-X scanhead, Andor iXon 897 EM-CCD camera and Andor laser combiner. All imaging with live cells was performed within incubation chambers at 37°C and 5% CO2. For RFP-Rab7, LAMP1-RFP and GFP-LC3 imaging, z stacks were acquired every 30 sec following addition of OPZ. Image acquisition and analysis was performed with Andor iQ3 (Andor Technology, UK) and ImageJ.

For analysis of GFP-LC3 recruitment to phagosomes, IFNg treated RAW264.7 cells pre-treated with inhibitors as indicated, followed by stimulation with OPZ for 25 min at 37°C. Z stacks from multiple fields of view were acquired by spinning disk confocal microscopy as described above. Maximum GFP intensity from line profiles made over multiple individual phagosomes from 3 independent experiments, were measured using Image J software.

For scoring cells with GFP-LC3 positive phagosomes, those cells containing 1 or more phagosomes were counted and scored positive if they contain any GFP-LC3 labelled phagosomes. More than 150 cells were analysed over 3 experiments.

For lysotracker imaging, IFNg treated RAW264.7 cells were incubated with LysoTracker Red (50 nM) for 15 min at 37°C. Z stacks were acquired over time, or after 25 mins stimulation with OPZ, by spinning disk microscopy as described above. Mean fluorescent intensity of individual phagosomes was measured using Image J software. For primary cells, Lysotracker Deep Red (50 nM) was used and images acquired using a Zeiss LSM 780 confocal microscope (Carl Zeiss Ltd), using Zen software (Carl Zeiss Ltd).

For entosis, MCF10A cells were plated on glass bottomed 6-well plates (MatTek). The next day, cells were maintained in an incubation chamber at 37°C and 5% CO2, and cell-in-cell structures imaged by widefield timelapse microscopy. Fluorescent and DIC images were acquired every 8 min using a Flash 4.0 v2 sCMOS camera (Hamamatsu, Japan), coupled to a Nikon Ti-E inverted microscope, using a 20 × 0.45 NA objective. Image acquisition and analysis was performed with Elements software (Nikon, Japan).

### Fixed immunofluorescent confocal microscopy and analysis

Cells were seeded in 12-well plates containing coverslips and incubated at 37°C, 5% CO2 for 24 h. Following treatments, cells were washed twice with ice cold PBS then incubated with 100% methanol at −20°C for 10 min. The cells were then washed twice with PBS and blocked with 0.5% BSA (Sigma, A7906) in PBS for 1 h at room temperature. The cells were incubated overnight at 4°C with the primary antibodies, then washed x3 in cold PBS. Fluorescent secondary antibodies were used at a 1:500 dilution in PBS + 0.5% BSA and were incubated with the cells for 1 h at room temperature. The cells were washed x3 in cold PBS prior to being incubated with DAPI for 10 min at room temperature and then mounted onto microscope slides with ProLong Gold anti-fade reagent (Invitrogen, P36930). Image acquisition was made using a Zeiss LSM 780 confocal microscope (Carl Zeiss Ltd), using Zen software (Carl Zeiss Ltd).

For primary cells, maximal ATP6V1A intensity was quantified at the phagosome membrane and divided by the mean intensity within the cell perimeter using Image J software. For each condition 10 phagosomes within 4 images were quantified and repeated in 3 independent experiments. For RAW264.7 cells, maximum intensity from line profiles made over individual phagosomes were measured from 3 independent experiments using Image J software.

### Salmonella infection

Infections were performed with wild-type and *DsopF Salmonella enterica* serovar Typhimurium SL1344 which are kind gifts from Dr. Feng Shao. Late-log bacterial cultures were used for infection during experiments as outlined previously (D’Costa et al., 2015). Briefly, GFP-LC3 expressing HCT116 cells were seeded on 1 cm^2^ glass coverslips in 24-well tissue culture plates 48 h before use at a density of 100,000 cells/coverslip. Bacteria were grown for 16 h at 37°C with shaking and then sub-cultured (1:33) in LB without antibiotics for 3 h. Post subculture, bacteria were pelleted at 10,000g for 2 min, resuspended and diluted 1:100 in PBS, pH 7.2, with calcium and magnesium, and added to cells for 10 min at 37°C. The cells were then washed 3 times with PBS with calcium and magnesium. Selection for intracellular bacteria was performed at 30 min post-infection using 100 μg/ml gentamicin. At 1 h post-infection, cells were fixed with 4% paraformaldehyde in PBS at 37°C for 10 min.

### Nitroblue tetrazolium (NBT)/formazan assay

RAW264.7 cells plated on cover slips were incubated with 0.2 mg/ml NBT for 10 min at 37°C. Where indicated, cells were also pre-incubated with inhibitors prior to addition of OPZ for 25 min. Samples were then fixed by ice cold methanol and dark formazan deposits detected by DIC imaging on a Zeiss LSM 780 confocal microscope (Carl Zeiss Ltd), using Zen software (Carl Zeiss Ltd).

### Luminol assay

RAW264.7 cells were seeded in 96-well white microwell plates (ThermFisher, 136101) at 1×10^5^ cells/well and incubated at 37°C, 5% CO2 for 24 h. Growth media was removed and cells were washed with PBS. Any pre-treatments were incubated at 37°C, and a final concentration of 0.32 U/ml of HRP (Sigma, P8375) and a final concentration of 125 nM of luminol (Sigma, A8511) diluted in DPBS (Dulbecco’s Phosphate Buffered Saline with Ca2+ and with Mg2+, Sigma, D8662), supplemented with 45% glucose solution (Sigma, G8769) and 7.5% sodium bicarbonate solution (Sigma, S8761), were added to the cells and incubated for the final 3 minutes of the pre-treatment incubation at 37°C. Stimuli were then added in triplicate and measurements were acquired immediately in the MicroLumatPlus LB 96V (Berthold technologies) at 37°C.

### Membrane fractionation

MCF10A cells were seeded on a 15 cm dish and cultured for 48 h. Cells were stimulated in suspension as indicated for 1 h. Input, cytosol and membrane fractions were isolated using the Mem-Per Plus Membrane Protein Extraction Kit (89842, Thermofisher) following product guidelines. Protein concentration was measured by BCA assay and equal amounts loaded onto polyacrylamide gels for SDS–PAGE analysis.

### Immunoprecipitation

One confluent 15 cm plate of HCT116 and RAW 264.7 cells was used per condition. Following stimulation, cells were washed twice with ice-cold PBS then lysed in 700 μl of lysis buffer (50 mM Tris-HCl (pH 7.5-7.6), 150 mM NaCl, 2 mM EDTA, 0.8% C12E9, and protease/phosphatase inhibitors) per plate, on ice, and incubated for 20 min at 4°C. Lysates were then centrifuged at 13,500 rpm for 10 min at 4°C and then rotated with 10 μl of pre-equilibrated protein A beads per sample (Pierce Protein A Magnetic Beads, ThermoFisher 88845, RA228548) for 1 h at 4°C. Following separation by magnet, lysates were then rotated with 25 μl of pre-equilibrated anti-FLAG^®^ M2 Magnetic Beads per sample (Millipore, M8823) for 2 h at 4°C. The lysates were washes five times in adjusted lysis buffer (50 mM Tris-HCl (pH 7.5-7.6), 150 mM NaCl, 2 mM EDTA, 0.1% C12E9, and 1% Triton X-100) by rotating for 10 min at 4°C. Samples were eluted by boiling at 95°C in 25 μl 2x SDS-buffer for 10 min.

### Western blotting

Cells were scraped into ice-cold RIPA buffer (150 mM NaCl, 50 mM Tris–HCl, pH 7.4, 1 mM EDTA, 1% Triton X-100 (Sigma, T8787), 0.1% SDS (Sigma, L3771), 0.5% sodium deoxycholate (Sigma, D6750) and lysed on ice for 10 min. Lysates were centrifuged for 10 min at 10,000 *g* at 4°C. Supernatants were then separated on 10% or 15% polyacrylamide SDS–PAGE gels and transferred to polyvinylidene difluoride membranes. Membranes were blocked in TBS-T supplemented with 5% BSA for 1 h at room temperature and incubated overnight at 4°C with primary antibodies diluted in blocking buffer. They were then incubated with a horseradish peroxidase-conjugated secondary antibody and proteins were detected using enhanced chemiluminescence (GE Healthcare Life Sciences, RPN2209). Densitometry analysis of bands was performed using Image J software.

### Statistical analysis

Statistical analysis was performed using Graph Pad Prism software. One-way ANOVA followed by Tukey multiple comparison test or unpaired t test were used as indicated in figure legends.

### Online supplemental material

Fig. S1 shows GFP-LC3A and RFP-LAMP1 dynamics during entosis. Fig. S2 shows LysoTracker intensity of phagosomes treated with BafA1 or DPI. Fig. S3 provides additional data on ATG16L1-V-ATPase interaction upon monensin treatment. Video 1 shows GFP-LC3A and Rab7-RFP dynamics during phagocytosis. Video 2 shows GFP-LC3A and LAMP1-RFP dynamics during phagocytosis.

## Supplementary Data

**Figure S1.**
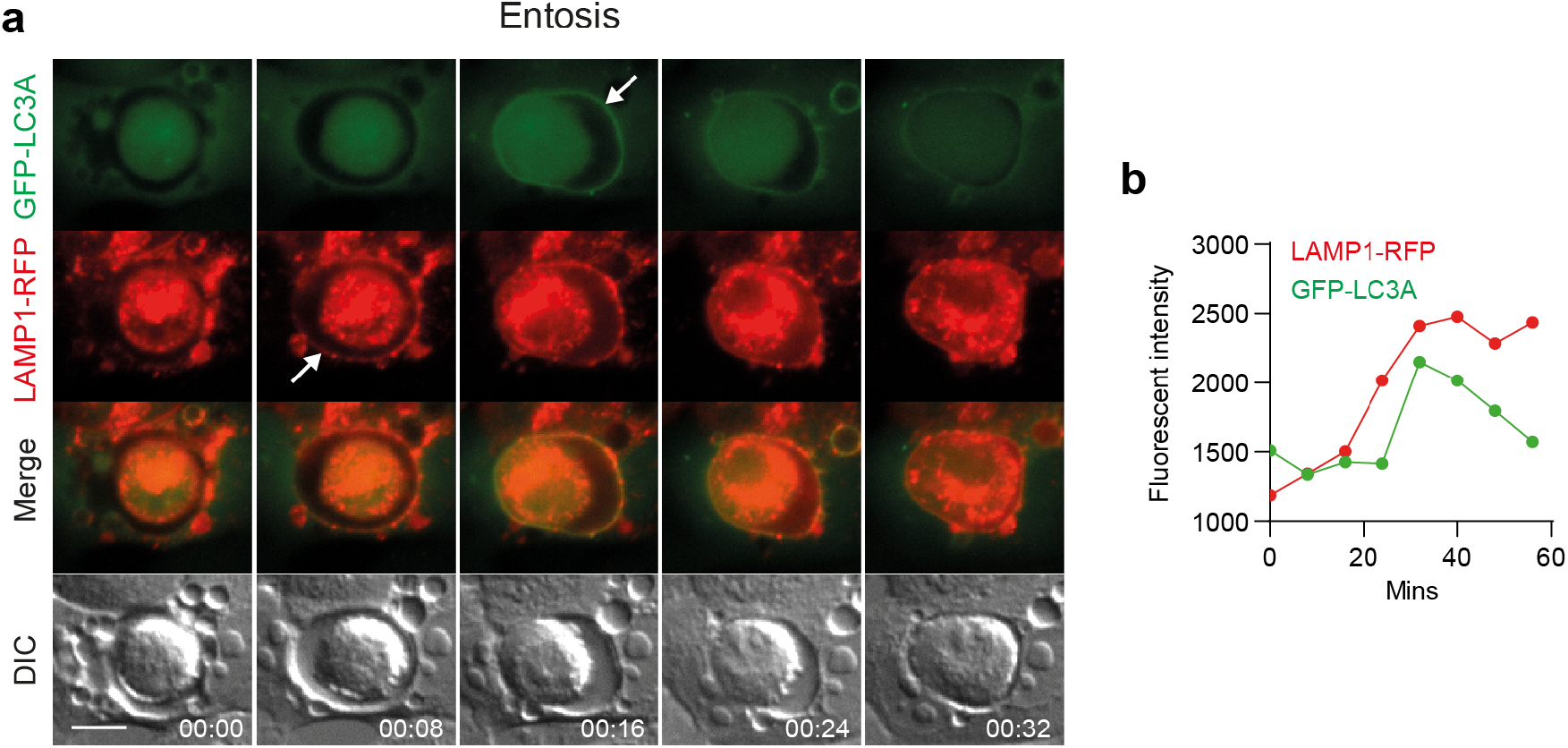
**(a)** Representative timelapse widefield microscopy images of entosis in MCF10A cells expressing GFP-hLC3A and LAMP1-RFP. Arrows mark acquisition of fluorescent markers. Scale bar, 15 μm, min:sec. **(b)** Analysis of fluorescent marker intensity at the entotic vacuole membrane over time.

**Figure S2.**
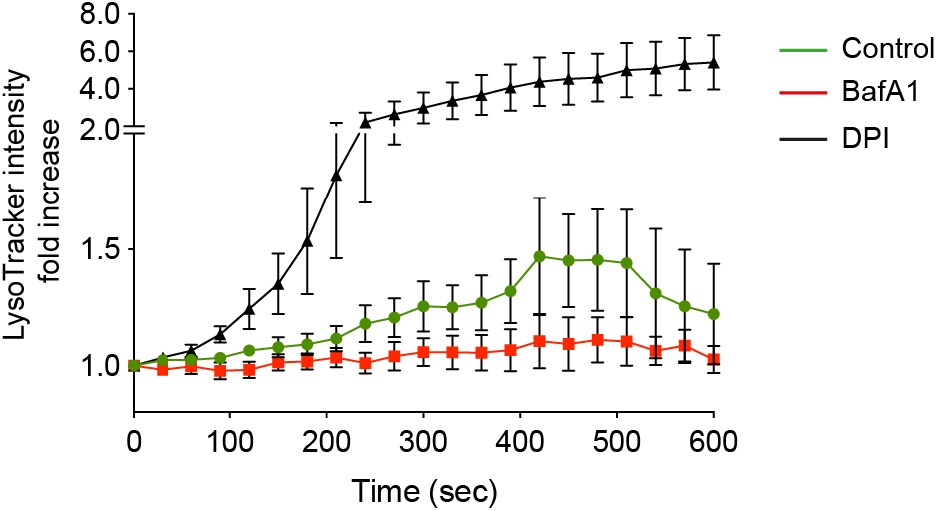
Confocal microscopy measurements of phagosome LysoTracker intensity over time in RAW264.7 cells pre-treated with either DPI (5 μM) or BafA1 (100 nM) prior to addition of OPZ. Data represent mean values normalised to time 0 +/-SEM of 9 individual phagosomes across multiple independent experiments.

**Figure S3.**
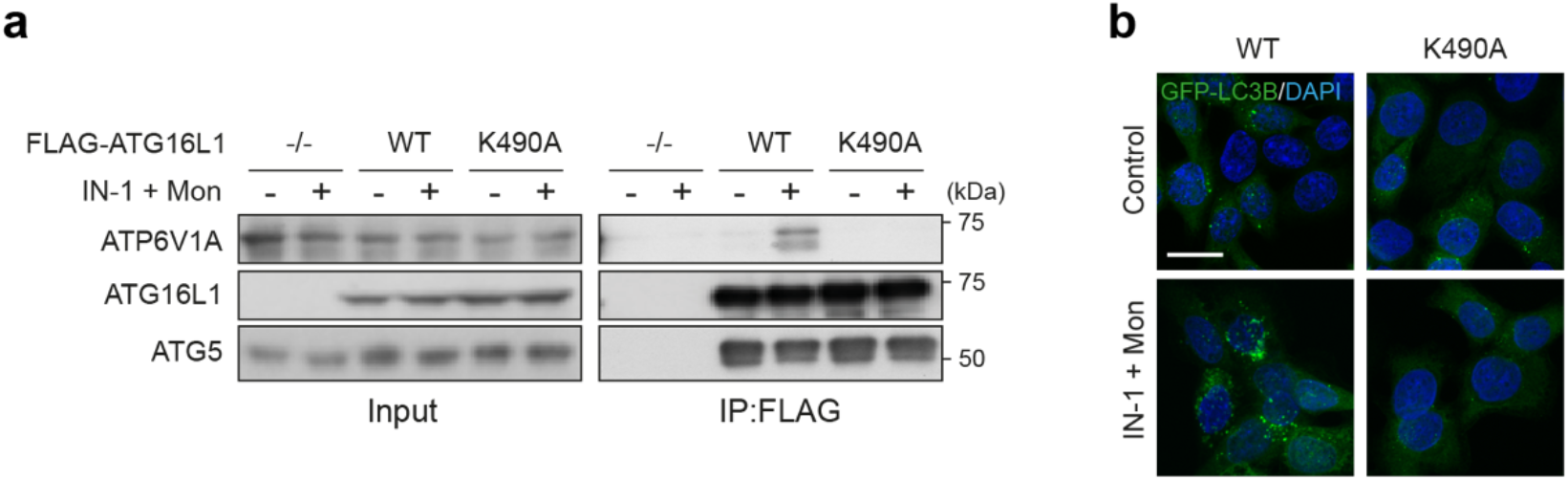
**(a)** *ATG16L1*^-/-^ HCT116 cells and those re-expressing FLAG-tagged wild type and K490A ATG16L1, were treated with monensin (100 μM) + IN-1 (1 μM) for 45 min. Input lysates and FLAG immunoprecipitations were probed for ATP6V1A, ATG16L1 and ATG5 by western blotting. **(b)** Confocal images of GFP-rLC3B HCT116 cells expressing wild type or K490A ATG16L1 treated as in (a). Scale bar, 15 μm.

**Video 1**

Timelapse confocal microscopy of opsonised zymosan (OPZ) phagocytosis in RAW264.7 cells expressing GFP-hLC3A and RFP-Rab7.

**Video 2**

Timelapse confocal microscopy of opsonised zymosan (OPZ) phagocytosis in RAW264.7 cells expressing GFP-hLC3A and LAMP1-RFP.

